# CFTR misfolds during native-centric simulations due to entropic penalties of native state formation

**DOI:** 10.1101/350074

**Authors:** J. Shi, T. Cleveland, A. C. Powe, G. McGaughey, V. S. Pande

## Abstract

Cystic fibrosis (CF) is a common genetic disorder that affects approximately 70,000 people worldwide. It is caused by mutation-induced defects in synthesis, folding, processing, or function of the Cystic Fibrosis Transmembrane conductance Regulator protein (CFTR), a chloride-selective ion channel required for the proper functioning of secretory epithelia in tissues such as the lung, pancreas, and skin. The most common cause of CF is the single-residue deletion of F508 (F508del), a mutation present in one or both alleles in 90% of patients that induces severe folding defects and results in greatly reduced expression of the protein. Despite its medical importance, high-resolution mechanistic information about CFTR folding is lacking. In this study, we used molecular dynamics simulation with a native-centric force field to examine the folding and assembly of both full-length CFTR and the isolated first nucleotide-binding domain (NBD1). We observed that the protein was capable of substantial misfolding on both the intradomain and interdomain scale due to entropically favorable kinetic traps that exist on CFTR’s folding free energy surface. These results suggest that even wild type CFTR, in the absence of any disease-related mutations, has suboptimal folding efficiency. We speculate that such entropically-driven misfolding also occurs in disease-prone mutants such as F508del and contributes to the protein’s poor *in vivo* activity.

## Introduction

Cystic fibrosis (CF) is a common genetic disorder that affects roughly 30,000 people in the United States and 70,000 worldwide (1). The disease is caused by certain mutations in the gene encoding the Cystic Fibrosis Transmembrane conductance Regulator protein (CFTR), a 1480 amino acid chloride ion channel expressed mainly in secretory epithelial tissue. The protein comprises two membrane-spanning domains (MSD1/MSD2) that form the channel pore, two nucleotide-binding domains (NBD1/NBD2) that are responsible for ATP-binding and channel gating, and a regulatory domain (R), which modulates gating in a phosphorylation-dependent fashion (2). Currently, there are more than 1700 confirmed disease-related mutations, with effects ranging from inducing synthesis of non-functional CFTR (class I), reducing folding and processing efficiency (class II), impairing channel gating (class III), reducing ion conduction through the channel pore (class IV), to reducing the quantity of functional CFTR synthesized in the cell (class V) (3).

The most common cause of CF is the class II deletion of F508 (F508del), which is present in one or both alleles in 90% of patients (1). This mutation causes severe folding defects and results in rapid degradation in the endoplamic reticulum (ER) and insufficient expression at the cell membrane (1). Previous experimental studies on F508del-CFTR suggest that the defective folding is due primarily to a combination of reduced thermodynamic stability of the NBD1 domain (4) and impaired domain-domain assembly of NBD1 with an intracellular loop, ICL4, present on MSD2 (5). Correcting these defects has been shown to be possible by reducing temperature, introducing rescue mutations to the NBD1 domain, or using small-molecule correctors (6,7,8,9). Considerable progress has already been made in the development of therapeutics for CF patients with F508del-CFTR (10). However, given that there is still neither a high-resolution mechanistic model of CFTR folding nor definitive structures of the misfolded conformations targeted for degradation in the ER, the process of rationally designing effective therapeutics remains a significant challenge.

In this study, we attempt to gain a mechanistic understanding of the folding and assembly of wild type CFTR using molecular dynamics simulation (MD). We employ a coarse-grained model in which only the Cα atoms of the backbone were explicitly modeled and use a Gō-type potential to model non-bonded interactions, which assigns an attractive force between pairs of natively contacting residues and a repulsive force between all other pairs (11). Such “native-centric” models have been successful at modeling the folding of many proteins (12). We do not explicitly model cellular machinery such as chaperones in our coarse-grained model, but instead make the simplistic assumption that the presence of chaperones places the protein in an environment conducive to folding, which is modeled by the native-centric potential. Utilizing such a coarse-grained approach was necessary for computationally modeling CFTR due to the long folding and assembly time, 30-120 minutes (13), which is several orders of magnitude longer than the limits of what is currently accessible if we were to represent the system on an all-atom scale using standard force fields (i.e. ~ milliseconds with current compute capabilities) (14).

Two separate sets of simulations were run: one for the full-length protein, and one for isolated NBD1 that additionally modeled the effects of co-translational folding. In both cases, we observed a significant amount of misfolding, which suggests that wild type CFTR has suboptimal folding efficiency both on the intradomain and interdomain scale. We hypothesize that these misfolding tendencies may be exacerbated in F508del-CFTR and contribute to the mutant’s poor *in vivo* expression. Our results provide experimentally testable structural insights into the misfolding of wild type CFTR that may potentially generalize to disease-related mutants.

## Results

### The domain-domain assembly efficiency of NBD1 with NBD2 is suboptimal

The putative native state used for the full-length CFTR native-centric simulations was a homology model of the Sav1866 ABC transporter protein created by Mormon and Callebaut (15). We first unfolded this structure at high temperature, and then initiated a total of 324 independent refolding trajectories, each of 5 × 10^4^ frames in length (see Methods). During this refolding process, we observed proper folding of CFTR to the native state in 63% of the trajectories, which occurred over three phases with distinct timescales. First, the MSDs and NBDs independently and rapidly folded into their native conformations, with the NBDs (NBD1: 3769 ± 272 frames, NBD2: 4148 ± 216 frames) folded more slowly than the MSDs (MSD1: 2795 ± 162 frames, MSD2: 2565 ± 181 frames). Then, assembly occurred between MSD1/NBD1 (6388 ± 413 frames), and MSD2/NBD2 (5432 ± 349 frames) to form two halves of the protein: MSD1/NBD1 and MSD2/NBD2. These halves of the protein are separated by more than 60 Å immediately after formation. Finally, these two halves diffused toward each other and assembled cooperatively into the native channel, with MSD1 assembling with MSD2 (14680 ± 497 frames) and NBD1 with NBD2 (14895 ± 517 frames).

In the remaining 37% of the trajectories, we observed misfolding and/or misassembly of the protein. In 43% of these remaining trajectories, we observed proper MSD1/MSD2 assembly but improper assembly of NBD1 and NBD2 (Figure 1, top). If the cores of both NBDs faced each other during assembly, MSD1/NBD1 and MSD2/NBD2 could rapidly assemble into the native channel. However, if at least one of the cores faced away from the other, this process was inhibited, resulting in the formation of long-lived conformations that exposed the NBD cores (Figure 1, top). This process is irreversible for the length of the simulation (5 × 10^4^ frames), which is approximately 3x the mean native folding and assembly time of ~1.5 × 10^4^ frames. This suggests that these misfolded states may be sufficiently long-lived to be targeted for degradation in the ER, which has a timescale of approximately 30 minutes (16) comparable to that of CFTR assembly (13). Alternatively, in 17% of these trajectories, both the MSDs and NBDs failed to assemble properly (Figure 1, bottom). In total, some form of interdomain misassembly occurs in 60% of the misfolding trajectories.

**Figure 1:**
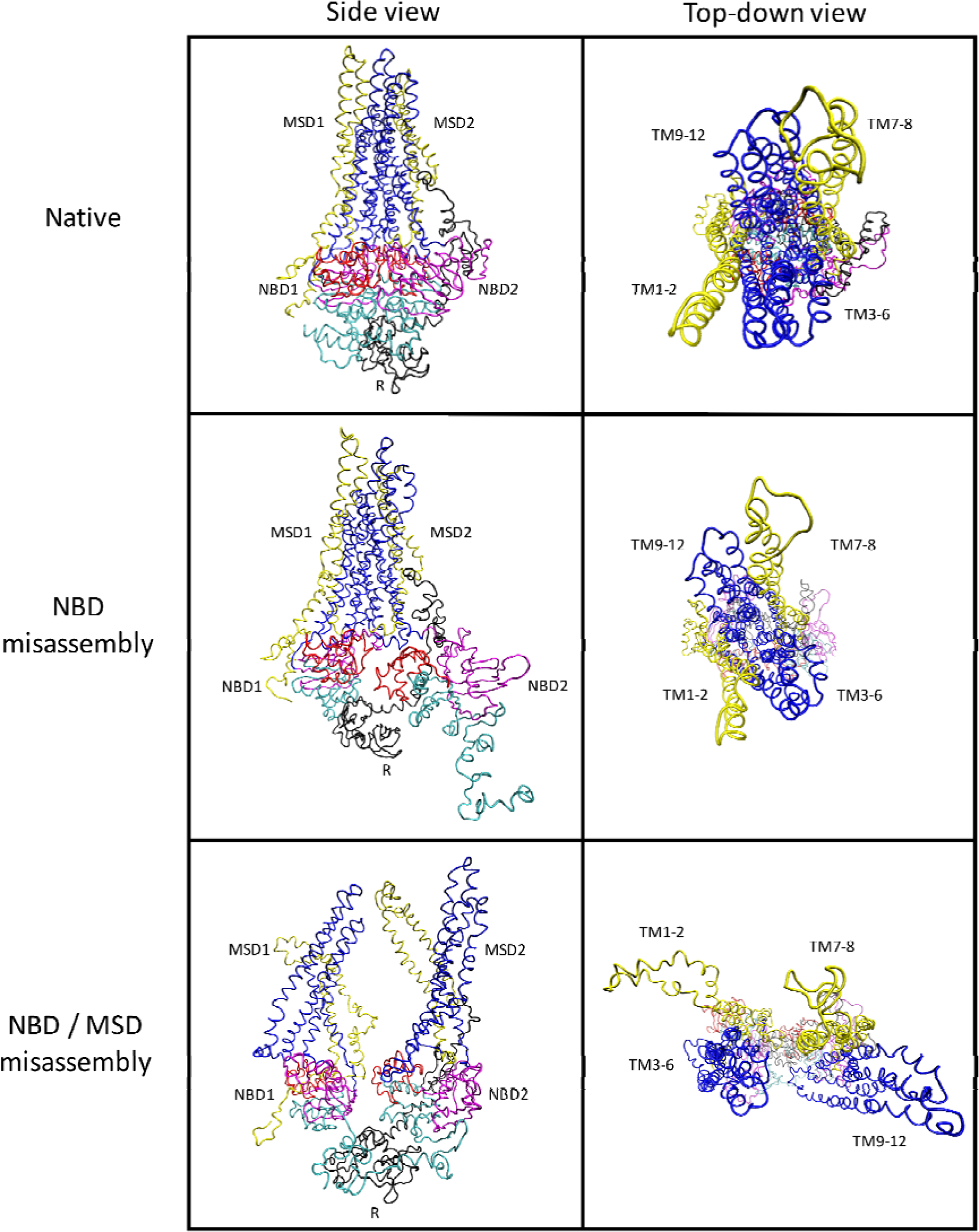
CFTR misassembles into two main species during the folding assembly process. The dominant species (middle) comprises 43% of the non-native trajectories and shows properly assembled MSDs but improperly assembled NBDs. The other species comprises 17% of the trajectories and show improper assembly for both the MSD’s and NBD’s (bottom). The misfolded structures shown represent the RMSD centroids of the last frames of the all the corresponding misfolding trajectories.

In the final 40% of the misfolding trajectories, we observed intradomain misfolding of either NBD1 or NBD2 into long-lived non-native species. We speculate that this is due to the large contact order between the N and C-terminal strands in the protein core, resulting in large entropic penalties for core formation. However, because our simulation does not model co-translational folding, the mechanism by which NBD1 folds *in vivo* (17,18), and instead models the domain as folding from a fully synthesized random coil, the folding dynamics obtained from our simulations are likely not physically realistic and we will therefore not discuss these species further. Rather, we decided to more closely investigate the mechanism of NBD folding by running additional folding simulations of isolated NBD1 under different translation conditions to see if the NBD folding inefficiency observed in the full-length simulations is preserved. These simulations are discussed in the following section.

A summary of the complete folding energy landscape seen in our full-length simulations is shown in Figure 2. Overall, we observe one dominant folding pathway comprising 63% of the trajectories, and three distinct, irreversible misfolding pathways, each resulting in the protein being trapped in a long-lived local energy minimum.

**Figure 2:**
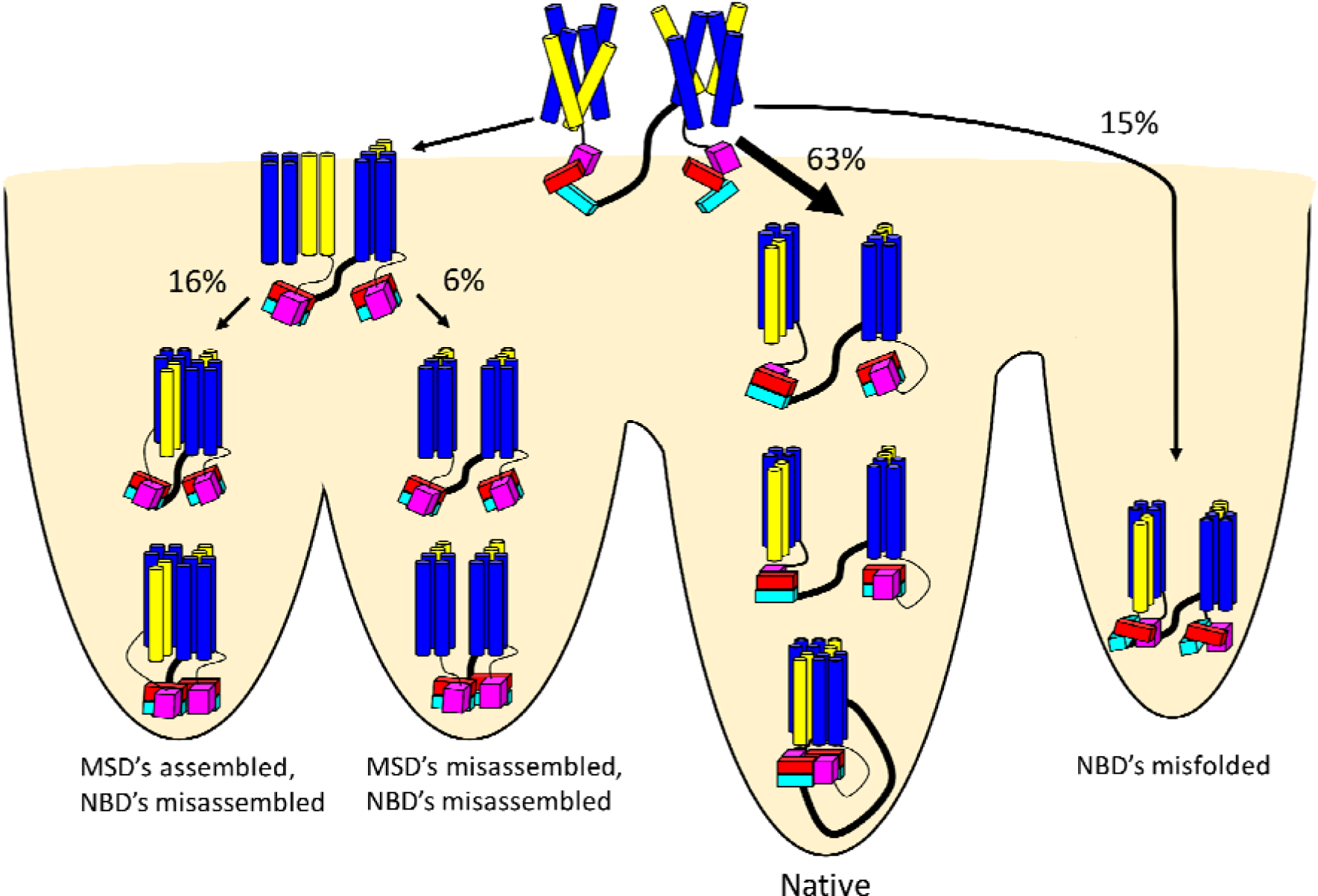
We observe four distinct mechanistic pathways for CFTR folding and assembly. The dominant process is the protein’s folding successfully into the native state. However, we also see a substantial degree of interdomain misassembly (left branches) as well as intradomain misfolding of the NBD’s (right branch).

### Modeling NBD1 co-translational folding reveals potential folding inefficiencies in the protein core

We observed a significant fraction of NBD misfolding in the full-length CFTR trajectories. Therefore, we decided to further investigate whether these misfolding tendencies were also observed when co-translational folding effects were included in the model. For this purpose, we simulated isolated NBD1 using two models that were designed to represent the limits of infinitely fast and slow ribosome translation relative to the rate of folding (hereafter referred to as fast and slow translation). We posited that the effects of *in vivo* translation should lie somewhere in between the results of these two models. We started these simulations with a pre-folded N-terminal subdomain, with the α-helical and C-terminal subdomains initialized in a straight chain. This choice of starting structure with a pre-folded N-terminal subdomain was motivated by experimental evidence suggesting that the N-terminal subdomain folds rapidly and independently of the other subdomains (17). To model co-translational folding effects, two sets of simulations were performed:

1. The α-helical subdomain was first folded with the C-terminal subdomain constrained in a straight chain conformation. After the α-helical subdomain completed folding into its native conformation, these constraints were removed and the C-terminal subdomain was allowed to fold. This model assumes that subdomain folding is infinitely fast compared to translation.
2. The α-helical and C-terminal subdomains were folded concurrently starting with a straight chain without constraints. This model assumes subdomain folding is infinitely slow compared to translation.

Attractive Lennard-Jones potentials were also added between all pairs of hydrophobic residues to model the effects of non-native hydrophobic collapse. 140 and 180 separate trajectories were run for models 1) and 2) respectively, each simulation lasting 5 × 10^3^ frames.

The most striking observation from these simulations is that under both modeled translation rates, the core of the protein, comprising strands S3 and S6 from the N-terminus and strands S7-S10 from the C-terminus, showed suboptimal folding efficiency. The intercalation efficiency of S8 into the N-terminal S6/S3 pocket is only 82 ± 3% for slow translation and 66 ± 4% for fast translation (Figure 3). The suboptimal intercalation was due to non-native hydrophobic compaction of the C-terminal subdomain, or packing against the α-helical subdomain, in either case inhibiting S8 intercalation into the core. Furthermore, there was a clear dependence of intercalation efficiency on translation rate, with fast translation leading to less efficient intercalation. This is likely due to an enhanced propensity of incompletely folded C-terminal subdomain (under fast translation) to pack against incompletely folded α-helical subdomain. We posit that this inhibits productive folding for both subdomains. To support this idea, we do see a corresponding drop in folding efficiency of the α-helical subdomain under fast translation (95 ± 2% for slow and 86 ± 3% for fast). Furthermore, unfolded α-helical subdomain provides more degrees of freedom for S8 to sample relative to intercalation, increasing the entropic penalty of intercalation. Finally, we observed that S9 intercalation was also suboptimal, but interestingly, unlike S8 intercalation, appeared independent of translation rate (slow: 72 ± 4%, fast: 69 ± 3%). Finally, the total folding efficiency to the NBD1 native state was noticeably affected by translation rate: 56 ± 4% for slow and 42 ± 4% for fast.

**Figure 3:**
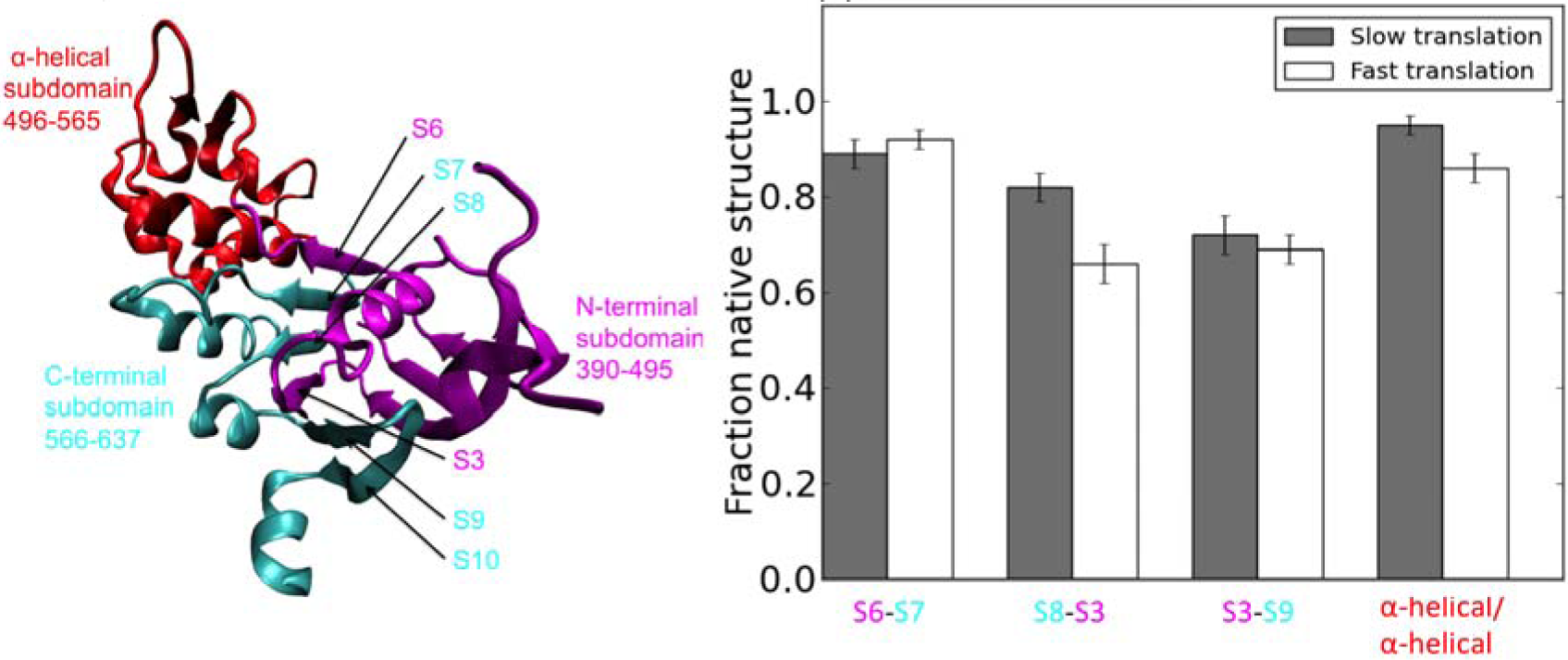
*Left:* NBD1 topology with subdomains color-coded: N-terminal (magenta), α-helical (red), and C-terminal (cyan). The core strands S3 and S6-10 are labeled. *Right:* Comparing the folding efficiency of the core of NBD1 under the fast translation model versus slow translation model at t = 5000 frames (~5x the average folding time of 1088 frames). Both fast and slow translation yields suboptimal folding efficiency for part of the protein core, i.e. S8-S3 and S3-S9 contact formation. The poorer S8-S3 contact formation under fast translation is due to hydrophobic collapse of S8 against an incompletely folded α-helical subdomain that both inhibits S8 intercalation and folding of the αhelical subdomain.

We hypothesize the suboptimal intercalation efficiency of S8 and S9 is due to the large separation in sequence, or high contact order (19), between the N-terminal and C-terminal strands of the central sheet. The hydrophobic compaction of the C-terminal subdomain and its ability to pack against the α-helical subdomain are entropically favorable competing pathways to intercalation into the core, leading to the formation of long-lived misfolded states. Interestingly, these results for wild type NBD1 are consistent with previous experimental studies of F508del that suggest the folding defect lies in the protein core (4) and may involve the last two strands (S9-S10) of the central sheet (20). Furthermore, the observed poorer intercalation efficiency of the core strands under fast translation is consistent with a study by Kim et al. that directly examined the co-translational folding of NBD1 at different translation speeds. In that study, it was found that increasing the translation speed, through either premature release from the ribosome or application of fast translation codons, resulted in reduced intercalation efficiency of C-terminal core strands and increased aggregation of the protein (18).

## Discussion

In this study, we set out to better understand the folding dynamics of CFTR using native-centric molecular dynamics simulations. We observe a common tendency for both full-length CFTR and isolated NBD1 to misassemble and misfold respectively even in the absence of any unfavorable mutations such as F508del. This suboptimal folding efficiency is caused by the large entropic barrier to both full-length CFTR assembly and NBD1 folding, and the availability of several off-pathway kinetic traps.

In the case of full length CFTR, we observed a substantial degree of misassembly of NBD1 and NBD2, and to a lesser extent MSD1 with MSD2. This is caused by the non-negligible thermal probability of these domains folding into misaligned configurations that inhibit assembly. The key assumption in our simulation is that the motions of the MSD1/NBD1 and MSD2/NBD2 portions of the protein are independent of each other, and that each portion has the freedom to face one of many different directions about the z-axis once it has completed folding on an intra-domain level. Essentially, we treat the direction faced by these halves of the protein as determined by two independent Bernoulli processes, with probability *p* of each half facing a suitable direction to allow proper assembly. Given our observed folding efficiency of 0.63, *p* is simply the square root of this number, or approximately 0.8. Thus, in our simulation, there is roughly an 80% chance for either portion of the protein to be suitably oriented for assembly post intra-domain folding. However, this may be an underestimate of the true entropic penalty to assembly, because our starting structures for refolding were generated by unfolding the native state, and there may be some residual memory of the native state topology in our starting structures that biases the protein towards correct assembly. Nevertheless, even under these circumstances, we still observe that the entropic penalty for assembly is sufficiently large such that a substantial fraction of the full-length trajectories still misassemble.

We must note that recently, Li et al. solved a cryo-EM structure for the phosphorylated and ATP-bound “open” configuration of the full-length human CFTR channel (21). Comparison of this structure to the “open” Sav1866 homology model used in these simulations revealed some clear structural differences between the structures (21). Specifically, the Sav1866 model assumed that the packing of MSD1 against MSD2 was symmetrical and exhibited identical domain swapping topologies for both the TM1-2/TM9-12 and TM7-8/TM3-6 interfaces. In contrast, the cryo-EM structure has TM7-8 packed against the four helices in its own domain, MSD2, rather than against the four helices in MSD1 per the domain-swapping hypothesis. This potentially calls into question the accuracy of our simulations, as the target native state upon which we based our native-centric force field contains structural differences from the cryo-EM structure.

However, we argue here that misassembly observed in our simulations should be robust to these structural differences. Firstly, we observe that although the TM7-8/TM3-6 interface in the cryo-EM structure is unique from that of the Sav1866 model, the interface between TM1-2/TM9-12 is well-preserved, and shows the domain-swapping topology predicted in the Sav1866 model. Due to this similarity, we posit that improper assembly of this interface can still occur even if we use the cryo-EM structure as the target native state. Furthermore, because TM7-8 in the cryo-EM structure is no longer packed against TM3-6, stabilizing native contacts between MSD1 and MSD2 involving these helices would no longer be included in the native-centric force field, leading to an overall fewer number of attractive potentials to stabilize the native state. As a result, a model built from the cryo-EM structure may even be *more* prone to misassembly due to the loss of these contacts. The NBD1/NBD2 interfaces are also essentially identical, so we predict that analogous misassembly of the NBDs could also occur.

In the case of isolated NBD1, we propose that the high contact order between the C-terminal and N-terminal core strands (146 residues for S8-S3 and 164 residues for S3-S9) results in slow formation of the core. From a contact order standpoint, non-native hydrophobic collapse of the C-terminal domain against the α-helical subdomain or compaction against itself are both entropically favorable competing mechanisms that inhibits intercalation of C-terminal strands into the core and traps the protein in long-lived misfolded conformations. This misfolding propensity of the core was also noticeably affected by the choice of translational model. Under our model for infinitely fast translation with respect to subdomain folding, we noticed a 16% decrease in intercalation efficiency compared to slow translation. An experimental study by Kim et al. showed that substituting fast-translating codons into the α-helical subdomain resulted in increased aggregation of the protein (18). Given our results, we propose that this aggregation may be due to hydrophobic packing between kinetically trapped NBD1 molecules consisting of stalled, incompletely folded α-helical and C-terminal subdomains. Similarly, Kim et al. also observed that premature release of NBD1 from the ribosome immediately after the α-helical subdomain fully emerges into the cytosol reduces the intercalation efficiency of the C-terminal strands into the core (18). This premature release rapidly exposes an additional group of approximately 40 C-terminal residues sequestered in the ribosome exit tunnel to the cytosol, comprising both S7 and S8 (18). We posit that rapid exposure of these residues to the cytosol results in the C-terminal domain folding concurrently with an incompletely folded α-helical subdomain that is still relatively disordered, similar to the initial conditions of our fast translation model. Therefore, we hypothesize that premature ribosome release may encourage formation of similar kinetic traps observed in our fast translation simulations via a similar mechanism.

Overall, our results provide insights into possible misfolding mechanisms of both full-length CFTR and the NBD1 domain that may result in suboptimal activity *in vivo.* However, due to the incredibly complex folding environment *in vivo* and the relative simplicity of our simulation model, it is critical that our results be subject to experimental validation. Specifically, it has been shown that the folding efficiency of CFTR when expressed endogenously in T84 and Calu3 cells is actually near-optimal (92%) (22), although substantially less efficient expression in which roughly 50% of synthesized CFTR is degraded has also been reported in the same cell lines (16). The fact that such high folding efficiency has been observed experimentally suggests that our simple native-centric model is not fully capturing the complex chaperone activity *in vivo*, thereby underestimating the true folding efficiency and presenting an inaccurate picture of CFTR folding and assembly dynamics. Therefore, to validate the misfolded species observed in the full-length simulations, we propose a cysteine cross-linking experiment using two contacting residues on the NBD1 and NBD2 cores, which can only come into contact if the domains are oriented correctly during assembly. This experiment can be run concurrently with a previous experiment monitoring crosslinking between MSD1 and MSD2 (23). A difference between the degrees of crosslinking observed in these experiments could be used to infer the existence of a non-native species with assembled MSD1/MSD2 and misassembled NBD1/NBD2, the dominant misfolded species observed in our simulation.

With respect to NBD1 folding, although the decrease in intercalation efficiency of C-terminal strands S7 and S8 into the core under fast translation (premature ribosome release) has already been demonstrated by Kim et al. (18), the intercalation efficiency of S9 and S10, which we predict is roughly independent of translation rate (Figure 3), has not been explicitly demonstrated. This can be tested by a fluorescence quenching experiment utilizing the tyrosine residues on S10. Measuring the fluorescence of Y625 or Y627 in the presence of a quencher at the N-terminal subdomain, such as on S3, would allow for detection of the S3-S9-S10 portion of the central sheet, as tyrosine fluorescence would be significantly higher if the strands are not intercalated. Performing this experiment using both fast and slow codons would provide insights into whether intercalation of the final two C-terminal strands are indeed translation-rate independent. Finally, repeating the above S9/S10 fluorescence experiment with F508del-NBD1 and comparing with the wild-type NBD1 would test whether S9/S10 core intercalation is noticeably affected by the F508del mutation, which has been suggested based on previous studies but not explicitly shown (4,20).

## Conclusion

Employing coarse-grained molecular dynamics with a native-centric force field, we simulated the folding and assembly of both full-length CFTR and isolated NBD1. We show that even in an environment conducive to folding, wild type CFTR still has a significant propensity to misassemble on the interdomain scale and misfold on the intradomain scale due to the high entropic barrier to both correct channel assembly and NBD folding. These entropic penalties may persist or be exacerbated in disease-related mutants such as F508del, contributing to the poor folding efficiency of the mutant. Therefore, targeting the wild-type misfolded species with small-molecule correctors may improve folding efficiency in disease-causing mutants.

## Methods

### -Molecular dynamics simulation

The C-α native-centric simulation for full-length CFTR was prepared using the SMOG server (11) and was run using GROMACS (24). The reference native state for the protein was a homology model from Mormon and Callebaut built from a crystal structure of the Sav1866 protein, which, like CFTR, is a member of the ATP-binding cassette (ABC) family of proteins (15). A custom restraining harmonic potential was added to the initial SMOG force field for all residues along the z-axis (to which the MSDs were aligned) to keep them vertically oriented during folding and assembly. This potential was defined using a GROMACS restraints (posre.itp) file in which a harmonic coefficient of 0.5 is set in the z-direction for all atoms. This potential was meant to simulate the constraining effects the ER lipid bilayer exerts on the MSD’s during assembly. We observed that without this restraining potential, the protein would often undergo unphysical dynamics, such as the MSD1/NBD1 and MSD2/NBD2 halves of the protein flipping 180° in the z-direction with respect to each other, which is not possible *in vivo* due to the presence of the ER lipid bilayer. The usage of a native-centric potential over a standard force field to model non-bonded interactions was primarily used to accelerate the folding dynamics due to CFTR’s prohibitively long folding and assembly time. This also has the potential benefit of modeling the effects of the omitted cellular machinery such as chaperones that presumably place the protein in an environment conducive to folding. The protein was first unfolded at 200K for 5 × 10^6^ timesteps, and then refolded for 5 × 10^7^ timesteps at 130K with randomized velocities sampled from the Boltzmann distribution. The final aggregate dataset was a set of 324 trajectories with frames saved every 10^3^ timesteps to give trajectories of length 5 × 10^4^ frames.

The NBD1 simulations were prepared by first creating a truncated version of the full-length CFTR PDB file that contained only residues 390-637 that represents isolated NBD1. The SMOG server was then used to generate the corresponding GROMACS topology file, using default force field parameters, C-α coarse-graining, and no periodic boundaries. A straight chain of these residues was then built in PyMol (25), and the N-terminal subdomain was subsequently folded with the remainder of the protein constrained to the straight chain conformation. Restraining the straight chain was done using a GROMACS restraints file in which a large restraining harmonic potential in the x, y, and z directions were defined for all relevant atoms. This was accomplished by setting the harmonic coefficients in the x, y, and z directions to 10 for the α-helical and C-terminal subdomains residues in the posre.itp restraints file. The resulting structure, with folded N-terminal subdomain and straight-chain α-helical and C-terminal subdomains, was used as the starting structure for the NBD1 folding simulations.

Lennard-Jones-like potentials were then defined between all pairs of hydrophobic residues during the NBD1 simulations to model non-native hydrophobic collapse and appended to the original GROMACS topology file. The equilibrium contact length of these interactions was taken to be the average of the hydrophobic contact lengths in the native state, 8 Å. The strength of each non-native interaction was set to a range of values: 0.2x to 1.0x the strength of a native contact as defined in the original force field, in intervals of 0.2x, i.e. setting the depth of the potential energy well to 0.2x to 1.0x that of a native contact.

For the slow and fast translation models, 140 and 180 respective trajectories were run for each value of the non-native contact strength. Total simulation time for both slow and fast models was 5 × 10^6^ timesteps, with frames saved every 10^3^ steps, resulting in trajectories of 5 × 10^3^ frames. The results from the 1.0x non-native contact strength simulations are the ones discussed in the Results.

Finally, we note that we have expressed all timescales in units of simulation frames instead of physical time because our native-centric force field contains nonbonded potential functions that are artificially biased to accelerate the folding process and not meant to accurately model true non-bonded interactions. Due to this fact, our simulation timescales do not correspond to units of real time, and we therefore cannot accurately estimate physical timescales for the observed processes. However, we can establish relative timescales between processes (e.g. intradomain folding of full-length CFTR is roughly an order of magnitude slower than interdomain assembly), which is sufficient for our purposes in this study.

### Analysis of MD trajectories

Analysis of the MD data was done using the MSMBuilder (26) and MDTraj (27) software. Visualization of trajectories and protein conformations, and generation of all protein images, was done using VMD 1.9.2 (28).

The folding efficiency of full-length CFTR was obtained by computing the RMSD of the last frame of each trajectory to the reference Sav1866 homology model, and all trajectories with last-frame RMSD difference less than 7 Å were defined as successful folding trajectories. We also visually compared the last frame of each trajectory to the reference native state and obtained the same result. The folding and assembly times for individual domains and interfaces reported in the first paragraph of the Results were computed as the mean number of frames needed for that domain or interface to form 70% of its native contacts. The folding efficiency of the structural features in NBD1 reported in Fig. 3 were obtained by computing the mean fraction of relevant native contacts formed in the last frames across all trajectories. The native contact cutoff distance was set to 13.5 Å for both full-length CFTR and NBD1. This value was used because it was the minimum value required to identify 100% of the native contacts given the Cα full-length CFTR native state structure. The overall folding efficiency to the NBD1 native state was calculated by taking the structure in the last frame of each trajectory and computing the absolute difference in contact length per native contact between this structure and the reference NBD1 native state. All structures with a per-contact length difference of less than 1 Å were defined as folded. These structures were then individually checked by visually comparing to the reference native state using VMD.

## Author Contributions

J. S. ran the simulations, analyzed the data and wrote the paper. T. C., A. C. P. Jr., and G. M. prepared the homology model for full-length CFTR from which the native centric model was built. V. S. P. supervised the project.

## Acknowledgements

This work was funded by grant No. 5U19AI10966204 from the National Institute of Health.

